# Signatures of glassy dynamics in highly ordered lipid bilayers with emergence of soft dynamic channels

**DOI:** 10.1101/2024.11.22.624930

**Authors:** Harini Sureshkumar, Sahithya S. Iyer, Atreyee Banerjee, Pratyush Poduval, Edward Lyman, Anand Srivastavava

## Abstract

Over the last few decades, extensive investigations on spatial and dynamic heterogeneity have been performed on carefully reconstituted biological lipid membranes. Characterising the molecular features in heterogeneous membranes is extremely challenging due to experimentally inaccessible time- and length-scales of these emergent systems. In this context, simulations can provide important insights into molecular-level interactions leading to membrane heterogeneity and associated functions. To that end, we use the non-affine displacement (NAD) framework (a concept borrowed from Physics of granular material) to faithfully capture molecular-scale local membrane order in simulated heterogeneous bilayers. In our latest application of NAD, we investigate the temperature-dependent spatial and temporal organisation on microseconds trajectories of liquid-ordered bilayer systems at all-atom resolution (DPPC/DOPC/CHOL: 0.55:0.15:0.30; 40 nm x 40 nm with a total of 5600 lipids and 2 million atoms). Lateral organisation in these large bilayer patches show noticeable dynamic heterogeneity despite their liquid-ordered nature. Moreover, our NAD analyses reveal soft fluid channels within the tightly packed membrane reminiscent of the classical two-component Kob-Andersen glass-forming binary mixture. Hence, we characterised these systems using classical glass physics markers for dynamic heterogeneities such as Overlap, Four-point Susceptibility, van Hove, and Intermediate Scattering functions to quantify the multiple time scales underlying the lipid dynamics. Our analyses reveal that highly ordered membrane systems can have glass-like dynamics with distinct soft fluid channels inside them. Biologically, these dynamic channels could act as conduits for facilitating molecular encounters for biological functions even in highly ordered phases such as lipid nanodomains and rafts.

**Significance:** Liquid-ordered lipid membrane offers a reliable representation of the cholesterol-rich outer leaflet of the plasma membrane. We show that dynamic heterogeneity is present even in highly ordered phases of lipid membrane, which arise at equilibrium merely from thermal fluctuations. Membranes in this state act as strong glass formers, and are hence less susceptible to temperature fluctuations. The dynamics is faster than actin or protein-mediated reorganisation and presence of passive but dynamic “soft channels” amidst the highly ordered lipid environment suggests a new exciting possibility that these channels could act as fast conduits for molecular encounters on the robust membrane surface.

## I. INTRODUCTION

Cellular membranes are quasi two dimensional fluids comprised of a complex mixture of lipids and proteins. In mammalian plasma membranes, 30-40 mole % of this lipid mixture is cholesterol, a uniquely important lipid whose chemistry and interaction with other lipids underlies a complex phase diagram. In model membranes containing cholesterol and at least two other lipids (usually saturated and unsaturated lipids), a region of liquid-liquid coexistence is observed that is referred to as the liquid-ordered (L_*o*_) and liquid disordered (L_*d*_) phases.^1–3^. Similar behaviour is also observed in vesicles extracted directly from living cell membranes, which contain the full chemical complexity of the plasma membrane with hundreds of different lipids and proteins^4^. The mixing thermodynamics of lipid mixtures and the preference of different proteins for distinct local lipid environments is exploited by cells to organize their surfaces into platforms for signalling and other functions^2,5^. The details of lipid phases — their molecular structure^1^, partitioning of proteins^6–8^, and the diffusion rates and dynamics of protein encounter^9^— are therefore important for understanding membrane organization and function.

The liquid-ordered phase is of particular importance in this context, because the outer leaflet of the mammalian plasma membrane is particularly enriched in cholesterol and sphingomyelin^10^. About ten years ago, simulations of model membranes reported by Sodt, et al. revealed that the liquid-ordered phase is heterogeneous at the molecular scale, containing nanoscopic domains of local hexagonal order interspersed with more disordered regions^11,12^. This same structure was later observed in a binary mixture^13^, and is also supported by more recent NMR experiments^14^.

The dynamics of the liquid-ordered phase inherits aspects of this heterogeneous lateral organization. On short timescales, the collective lipid dynamics are well described by several low energy phonon modes, including a gapped optical mode resulting from the local hexagonal packing within the L_*o*_ phase^15^. On longer (diffusional) timescales, the L_*o*_ phase possesses an extended subdiffusive regime, observed in both simulations^16–18^ and ultra high speed single particle tracking^19^. Recent studies show that the diffusion rate itself follows a specific distribution, due to evolving length and time scales in glassy systems^17,20,21^. The anomalous diffusive dynamics, along with a striking visual similarity to configurations of polydisperse disks in 2D^22^, led us to revisit the dynamics of the L_*o*_ phase using concepts and approaches developed in the glass field,^23–27^ particularly an analysis of particle displacements in terms a “non-affine” parameter. Several of us have published a series of papers recently showing that this provides new insight into the complex structure and dynamics of the L_*o*_ phase^8,28–33^.

In this work, we use a combination of order parameters to quantify the time and lengthscales associated with spatial and dynamic heterogeneities within the L_*o*_ phase.. We also capture the emergence of “soft channels” in between the hexagonally ordered nanoscopic domains at near physiological temperatures. We reveal a non-trivial temperature dependence in the correlated length scale of soft channels using intermediate scattering function, overlap, dynamic susceptibility and van-Hove function. Using a non-affine order-parameter (χ^2^) analysis we also observe that the dynamic heterogeneity strongly correlates with variations in the local order across all temperatures and timescales. Originally introduced by Falk and Langer to study the viscoplastic deformation in amorphous solids^34^, χ^2^ has been recently shown to not only classify L_*o*_/L_*d*_ phases but also faithfully capture fluctuations in the phase boundaries^35–37^. We finally demonstrate that the χ^2^ non-affine parameter is strongly tied to the anomalous nature of lipid diffusion, making it ideal to study the heterogeneity of lipid systems, invariant of time, temperature and length-scales.

## II. SYSTEMS STUDIED AND SIMULATION DETAILS

This study involves investigations of lipid dynamics and local order in the ternary DPPC/DOPC/Chol lipid system near L_*o*_ phase. The initial configuration for the ternary L_*o*_ system consists of DPPC, DOPC and Chol lipids set in the ratio of 0.55:0.15:0.3 to recapitulate a predominantly ordered L_*o*_ phase according to the ternary phase diagram^38^ as shown in Fig. 1. The system consists of a total of 1.97 million atoms with 5600 lipids, 2400 cholesterols and 3,60,000 water molecules resulting in a system size of 40.8 nm×40.8 nm × 11.2 nm.

**FIG. 1.**
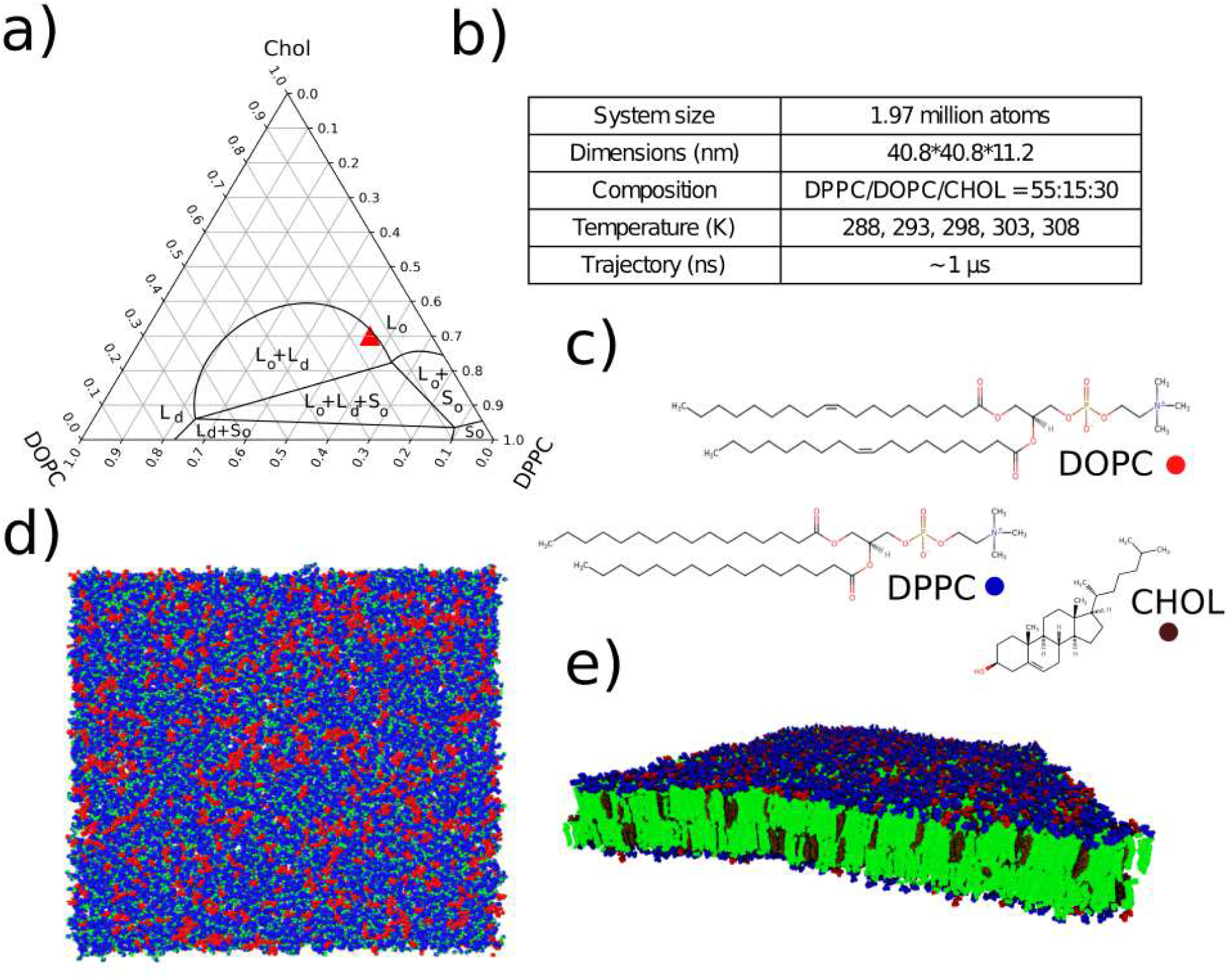
(a) Ternary phase diagram of DOPC-DPPC-CHOL membranes^39^ with composition of our system highlighted in the map (Red triangle); (b) details of the system under study; (c) molecular structure of DOPC, DPPC and Cholesterol generated using MarvinJS Chemaxon; (d-e) front and top view of the membrane patch color coded according to the lipid head groups (Red, Blue), tails (Green) and cholesterol (Brown)

The system was simulated at temperatures of 288 K, 293 K, 298 K, 303 K and 308 K in NPT ensemble using GROMACS 2021.3^40^. The temperature was controlled using Nose-Hoover thermostat^41,42^and semi-isotropic pressure coupling to set the pressure to 1 bar was achieved using Parrinello-Rahman barostat^43^. Each system was simulated for a minimum of 1 μs. The particle positions were collected every 10 ps for the analysis presented in next section. Input files for all five systems needed to initiate molecular simulations and generate the trajectory using Gromacs can be accessed via our laboratory GitHub link: codesrivastavalab/glassyDynamicsLipids. The trajectory files for all five systems under consideration can also be accessed directly from our SharePoint location, which is provided in the above github link and also on the following Figshare repository https://figshare.com/s/3adebead8c5785323449.

## III. CHARACTERISING DYNAMIC HETEROGENEITY ASSOCIATED WITH CHANNEL FORMATION

One of the hallmarks of glassy dynamics is the emergence of a distinct sub-diffusive regime in the mean squared displacement (MSD) profile given by 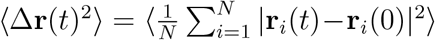 where N=number of particles and **r**(t) denotes the position at time t. The anomalous diffusion is tracked by its diffusion exponent (α(t)) given by,

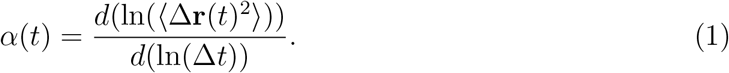

Fig.2(a) highlights a linear temperature-dependent lengthening of an already extended sub-diffusive regime (α(t)<1)^44^ in MSD, which is expected of a densely packed bilayer. The exponent eventually approaches Fickian diffusion (α(t)≈1) at microsecond timescales. The MSD graph is shown in the inset of Fig. 2(a). The diffusion exponent plateaus align with the first peak of radial distribution function *g(r)* (Fig. 2(b)), signifying a confinement radius of roughly 5 Å. Small neighbourhood implies tight packing of lipids for the ordered system under investigation. However the caging dynamics is dependent on temperature and timescales. The particle displacement profiles (Fig. 2(c-d)) for temperatures 288K and 308K, respectively show clear particle segregation that emerge at different timescales. We refer to them as ‘soft channels’ due to their dynamic fluid-like organisation in midst of packed order bilayer environment. The displacement organization is reminiscent of configurations observed in the classical Kob-Andersen like 2D binary mixture systems^22,45,46^. While Δr(t)^2^ and *g(r)* determine the caging regime, they fail in capturing the spatially and temporally localised fluctuations. Hence we map the dynamic heterogeneity using well-established measures in glass physics such as self-intermediate scattering function, overlap function, dynamic susceptibility and van-Hove function.

**FIG. 2.**
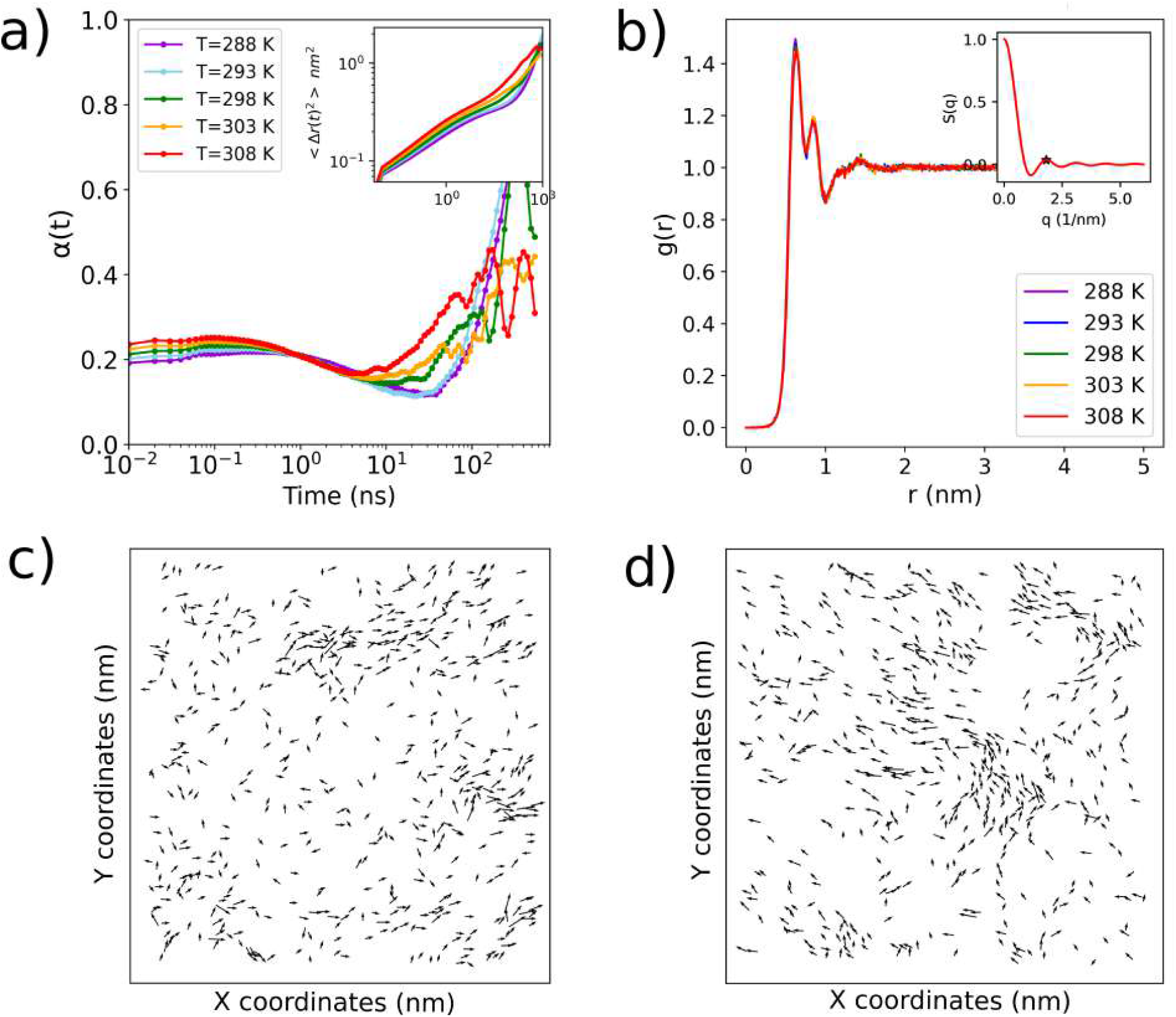
a) Exponent (*α*(t)) of mean square displacement function given in the inset b) radial distribution function (g(r)) and 2D structure factor S(q) (inset).The first peak of g(r) corresponds to 0.634 nm which is in agreement with S(q)=1.8 *nm*^*−*1^. Particle displacement vectors at temperatures 288 K (c) and 308 K (d) with a time lag of 260 ns and 60 ns respectively. Only particles with largest displacements (top 30%) are plotted here

### Self Intermediate scattering function

To measure the structural relaxation, we calculate the self intermediate scattering function F_*s*_(q, t) which is another universal feature of dynamical heterogeneity. F_*s*_(q, t) is given by,

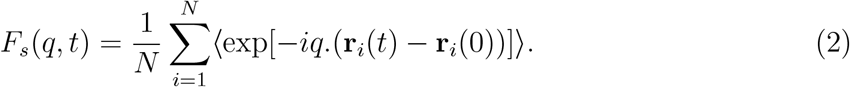

where the wavelength is give as 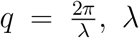, λ. To obtain the wave-vector dependence on the relaxation times, F_*s*_(q, t) are calculated at different values of q ranging from 1 to 3 nm^*−*1^. This range allows us to probe relaxation dynamics around the first peak of structure factor S(q) (Fig. 2(b) (inset)) which also corresponds to the length scale of caging. At low temperatures, the F_*s*_(q, t) transitions through two distinct temporal regimes: an initial phase of rapid ballistic motion within a confined “cage,” followed by cage relaxation processes and a subsequent slow escape from the cage.

In a homogenous system, the particles escape the cage evenly, leading to a smooth relaxation curve of exponential nature. The cage size is pre-determined by the wave factor q. In heterogenous systems such as ours, the particles exit the cages at various timescales, resulting in appearance of a secondary shoulder or plateau at intermediate timescales. At longer timescales all particles eventually decorrelate from its initial state indicating the start of normal diffusion (F_*s*_(q, t)=0). The F_*s*_(q, t) profile of the membrane undergoes a two-step relaxation process when the temperature is lowered. This is indicative of a growing effect of caging dynamics on the fast localised collective motions (q=2-3 nm^*−*1^) thus resulting in heterogeneous relaxation (Fig. 3). However, the large scale collective dynamics (q=1-2 nm^*−*1^) relax in a near exponential manner at all temperatures.

**FIG. 3.**
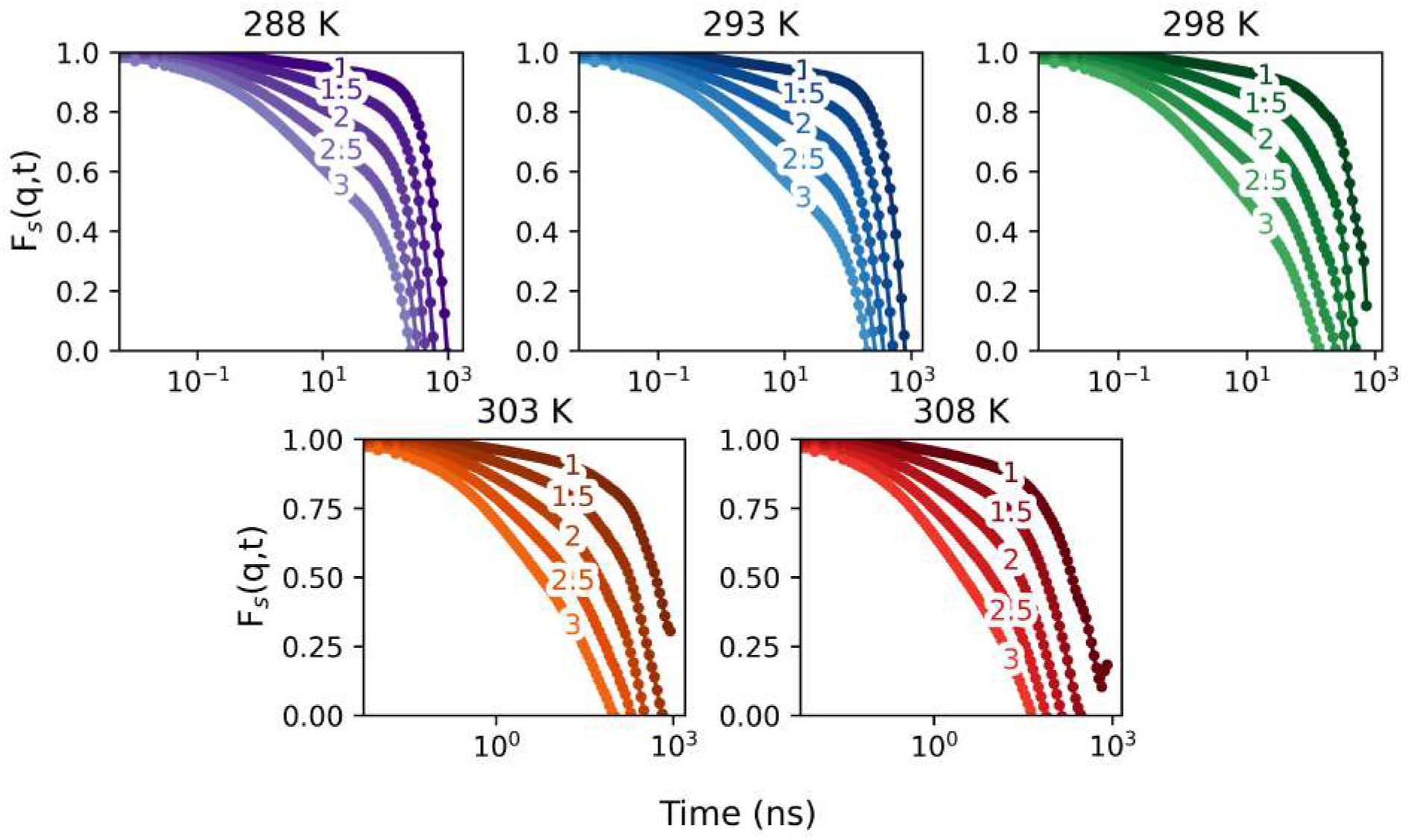
Self-Intermediate scattering function at q=1,1.5,2,2.5,3 at different temperatures.

### Overlap function

We now probe the short-time fluctuations in the caging region at distances corresponding to q=1.8 nm^*−*1^ using overlap function Q(t). The self-part of the overlap function Q_*s*_(t) gives the extent of correlation between two configurations separated by time *t*. It measures the number of particles that remains within a given distance (denoted as a for convenience here) of its initial position (at t = 0) after they evolve for time *t*. The Overlap function is expressed as,

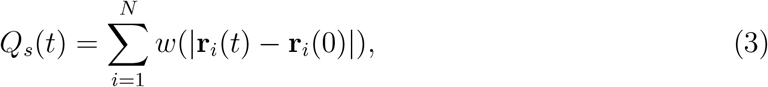

where w(|**r**_1_ − **r**_2_|) is 1 for particle separations less than or equal to distance a and 0 otherwise. We choose a=0.5 nm (≈ 1.8 nm^*−*1^) to coarsen out the amplitude due to vibrational motions prior to the caging phenomena.

The relaxation dynamics (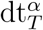 and 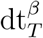) of Q_*s*_(t) signifies the timescales of correlations in lipid motions. At all temperatures, the lipid displacements decorrelate completely (Fig.4.a). This cross-over duration where the system decouples from its initial configuration and adopts Brownian motion is called alpha-relaxation (τ_*α*_). Physically, it is the time point (*dt*) where the function Q_*s*_(t) decays to 1/e of its initial value. The relaxation rates (τ ^*α*^) exhibit Arrhenius-like temperature-dependence. The relation is defined as 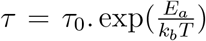 where τ = τ ^*α*^, T is the temperature and E_*a*_ is the activation energy. The activation energy barrier for the particles to enter diffusion regime is E_*α*_=62 kJ/mol which is comparable to the model membrane experimental estimates^47–49^. This observation indicates pronounced glass-forming characteristics in the membranes within this temperature range. Such behaviour may contribute to the maintenance of global structural integrity while simultaneously allowing localized relaxations that facilitate the formation of channels for signalling molecules.^50^.

The onset of localised relaxations, when particles escape confinement, is referred to as the β-relaxation regime (τ_*β*_). This presents in the overlap function as a lag in the exponential decay trend through the emergence of an intermediate secondary plateau or ‘shoulder’. τ_*β*_ is extracted by fitting a stretched exponent function - Kohlrausch-Williams-Watts (KWW) function of the form 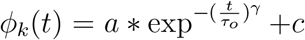, where τ_0_=τ_*β*_^51^ to the overlap function. However, the plateau in this system is not pronounced enough to determine the exponent γ for an accurate fit and extract τ_*β*_. Hence, we get an approximation of τ_*β*_ through dynamic susceptibility while simultaneously arriving at the KWW exponent (γ).

### Dynamic Susceptibility

Dynamic susceptibility (χ_4_) is a four point correlation function over particle displacements that quantifies the number of particles involved in correlated motions at time *t*,

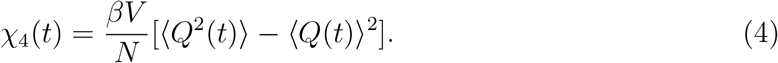

The fluctuation or variance of the Q(t) provides dynamic susceptibility. The value of these fluctuations in real space is a quantification of the lifespan and the size of transiently localized channels on the leaflet. The peak of χ_4_(t=τ_*peak*_) signifies maximally correlated motions in the system and subsequently the timescales of maximum heterogeneity. τ_*peak*_ grows and shifts to larger times indicating increasing length scales of DH (Fig.4.b). Since βrelaxation closely precedes τ_*peak*_, we approximate τ_*β*_ ≤ τ_*peak*_ and plug it in ϕ_*k*_(t) to quantify the extent of dynamic heterogeneity in the membrane. The γ values thus obtained, is consistently low (γ_288_=0.15 to γ_308_ =0.48), indicating multiple relaxation modes gradually vanishing with higher temperatures.

**FIG. 4.**
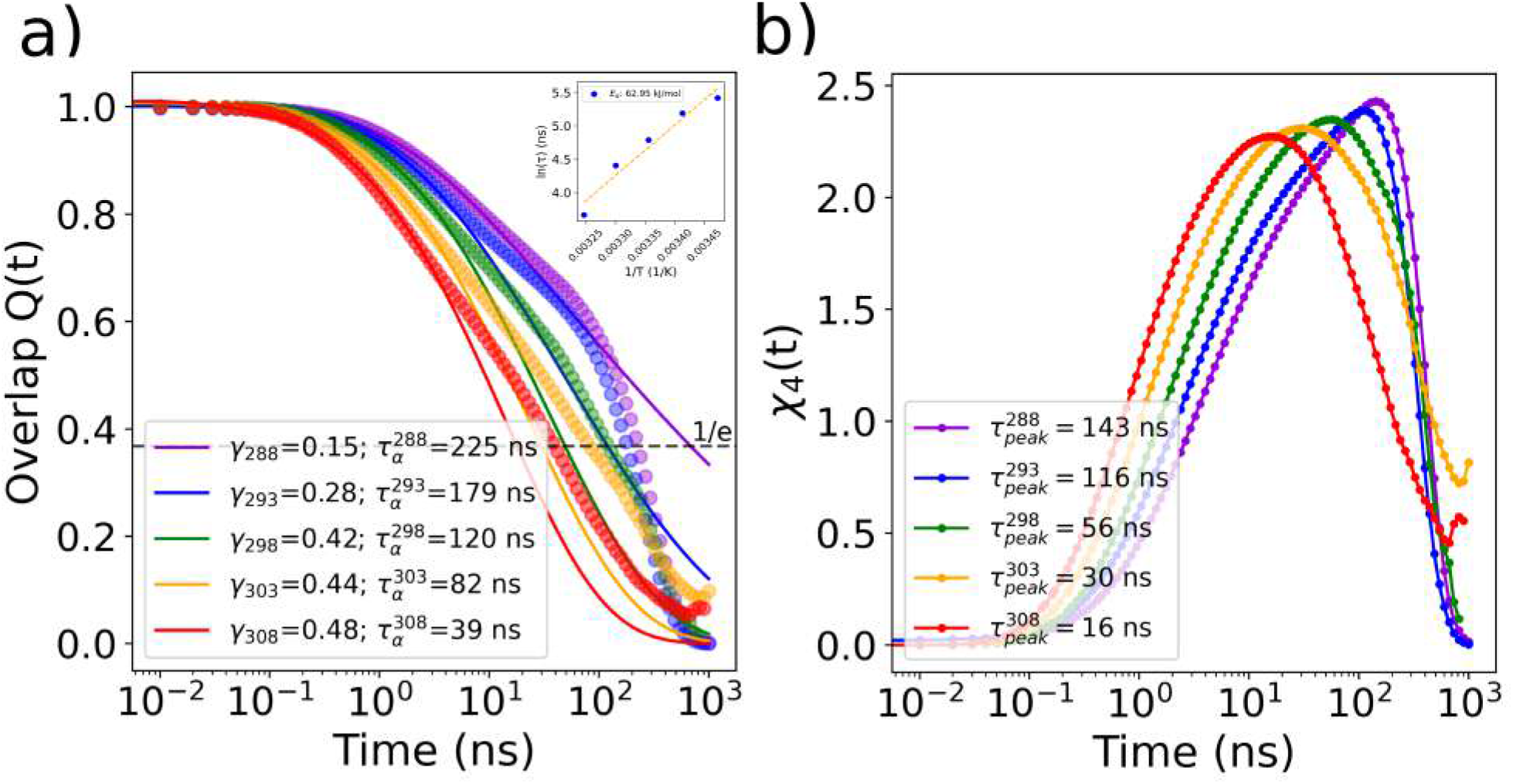
(a) Overlap function Q(t) for different temperatures (T). Legend contains values of exponent (*γ*_*T*_) fit to short timescales of Q(t) and 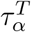 extrapolated at 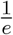 of its initial value (Inset) Arrhenius like behaviour of *α*-relaxation timescales with an activation energy of 62.95 kJ/mol depicted using a linear fit to ln(*τ*_*α*_) vs 1/T (b) Dynamic susceptibility *χ*_4_(t) across temperatures. The curve peak *τ*_*peak*_ is reported in legend.

We also visually observe a clear growing correlation length in particle displacements that sets in at timescales (τ_*β*_) slightly lesser than τ_*peak*_ and settles into stable channel forms postτ_*α*_. Please see the movie files provided as T288-Relaxation.gif for 288K in SI. Movie files for all other temperatures are also provided in the SI. Interestingly τ_*β*_ and γ deviate at 298 K. The cross-over between relaxation dynamics in F_*s*_(q,t) profiles for intermediate q, coupled with instability in γ fit indicate a shift in the underlying relaxation mechanism below 298 K. This is suggestive of a dynamic transition in the local packing order at this temperature, which is located on the tie-line between L_*o*_ and L_*o*_-L_*d*_ region in the phase diagram^39,52^.

### van-Hove function

To understand the spatial changes evolving with relaxation, we use van-Hove function to characterise spatial distribution of particles over time and apply nonGaussian parameter to quantify the deviation of particle distributions from the Gaussian behaviour.

The self part of van-Hove function G_*s*_(r, t) is a time dependent generalization of static density-density correlation function, also called the radial distribution function,

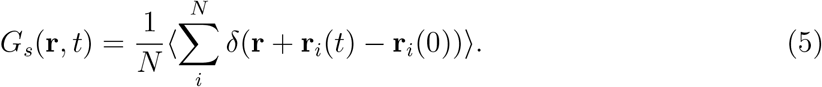

The probability distribution of particle displacements (2π **r** G_*s*_(**r**, t)) as seen in Fig. 5 is non-Gaussian at short timescales (dt<100 ns) for all temperatures and approaches Gaussian behaviour as timescales increase (Fig 5.f). In this region, the lipids are well-ordered and restrained by caging effects. At the lowest temperatures, there is a minor secondary peak (≈0.9 nm) for extremely short timescales (dt < 70 ns), indicating heterogeneous diffusion as some lipids intermittently escape their cages (0.634 nm: dashed grey line). This behaviour gradually decreases with increase in temperature and entirely vanishes above the dynamic transition temperature (298 K). This transition marks the cross-over from super-cooled heterogeneous dynamics to Brownian-like motion at 298 K with a characteristic timescale of 5 ns.

**FIG. 5.**
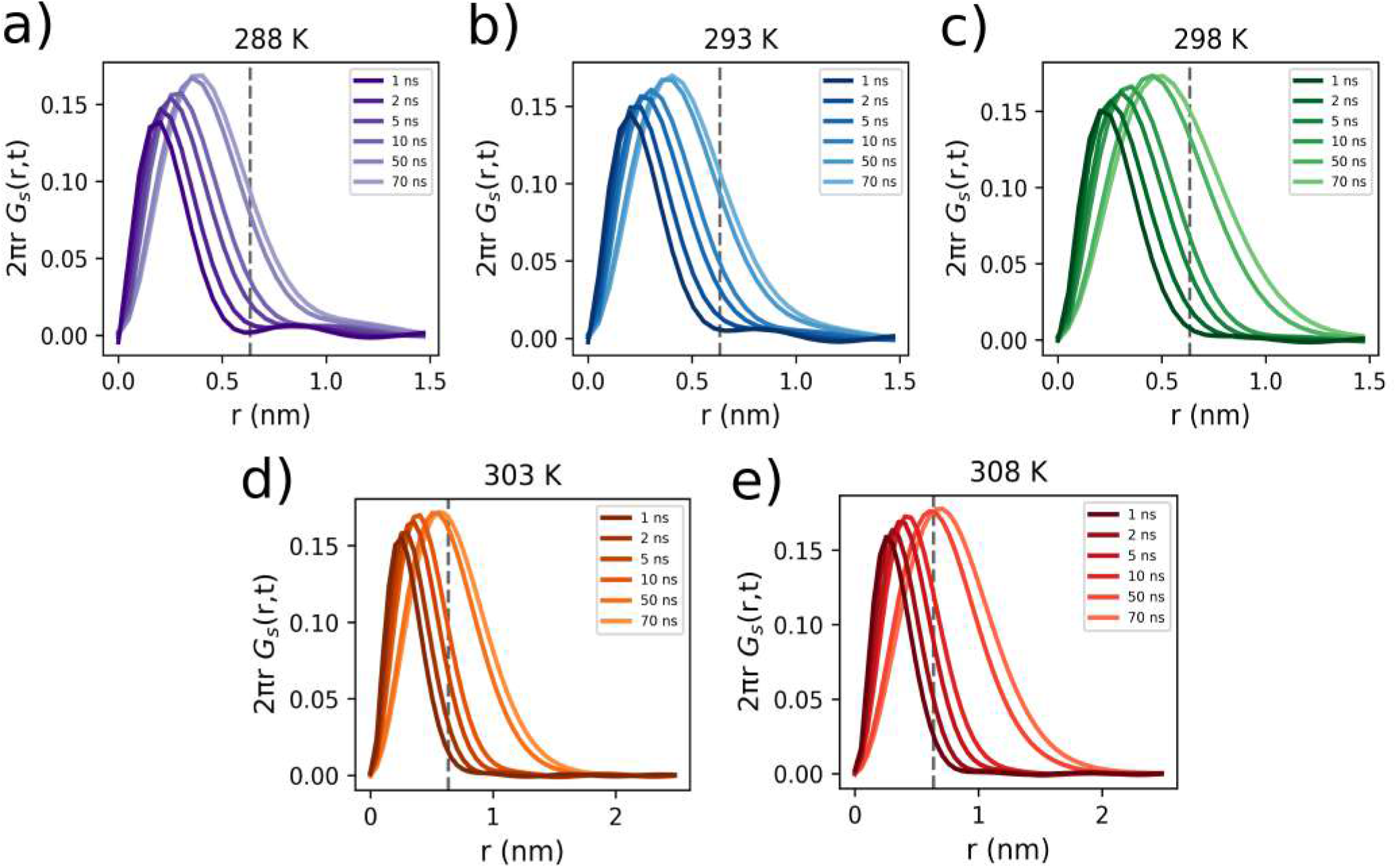
(a-e) van-Hove function evolving at different timescales (dt) for each temperature. Both the scales in this figure vary across temperatures to highlight the differences in distribution which is not visible otherwise, with uniform axis limits. The dashed grey line indicates the length of caging regime *λ*=0.634 nm

### Non-Gaussian Parameter

The deviation of van Hove distribution from Gaussian statistics is quantified using the Non-Gaussian parameter (α_2_(t)) as seen in Fig.6,

**FIG. 6.**
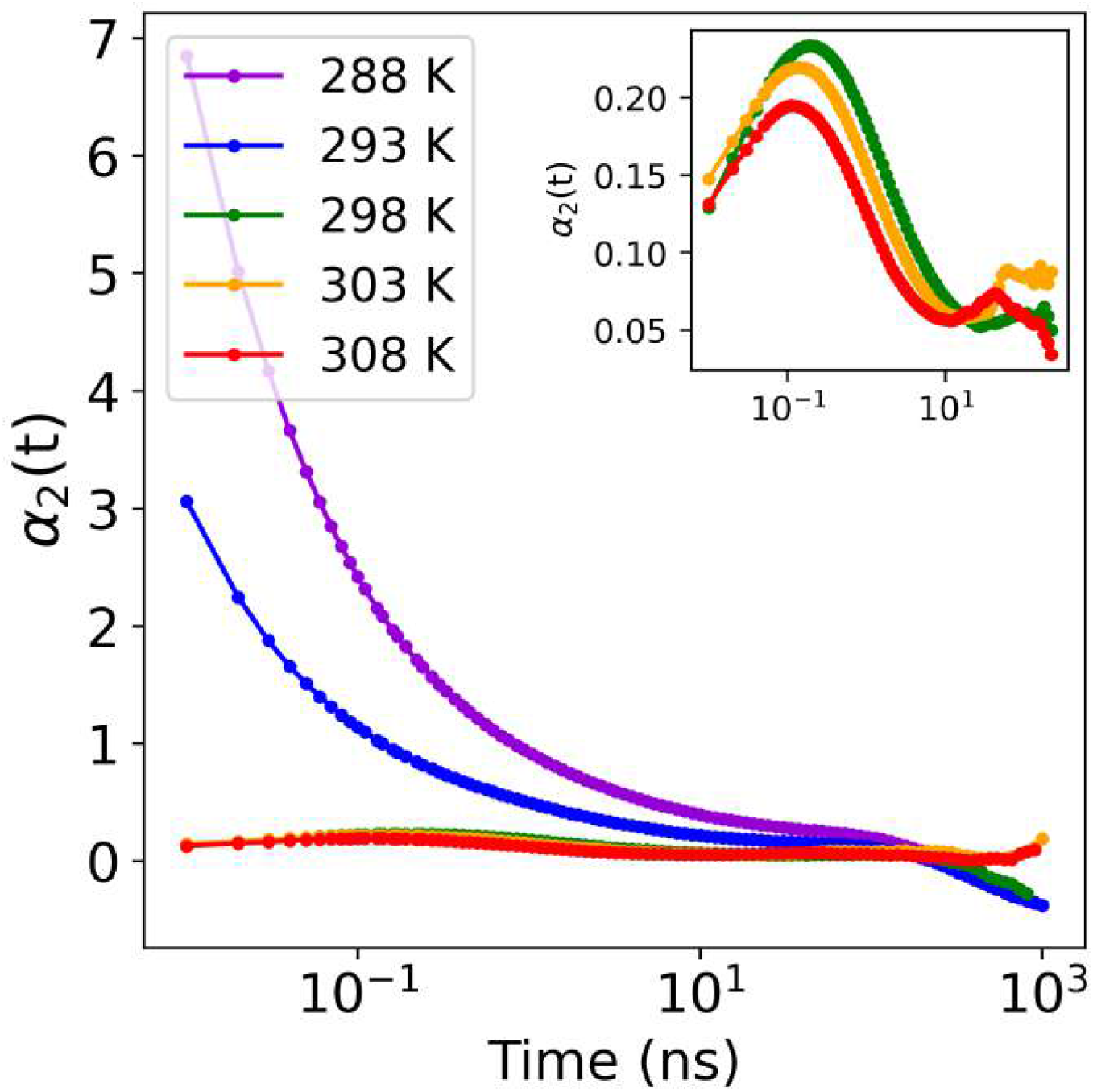
Non-Gaussian Parameter for each temperature. Inset: detailed profile of NGP at 298 K, 303 K and 308 K.

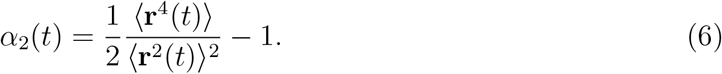

At higher temperatures, α_2_(t) is almost zero indicating uniform distribution of particles and Gaussian behaviour at all timescales. Below 298 K, the particle distributions are highly non-Gaussian thus validating the presence of mid-length correlations seen in Van-Hove at timescales less than 50 ns. It subsequently relaxes into Gaussian behaviour at 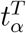 in concordance with relaxation dynamics of Q_*s*_(t). The increasing tail lengths with decreasing temperatures indicate extended timescales of correlated particle motions.

## IV. VISUALISATION OF DYNAMIC SOFT CHANNELS IN ORDERED MEMBRANES

The spatial map of displacement field (Fig.2(c-d) confirms the presence of transient localised clusters at different timescales identified by the dynamic heterogeneity analyses. Most active biological matter that display heterogeneities often pack in an intermediate hexatic phase, which retains both solid- and liquid-like properties. Hence, we employ hexatic order parameter (Ψ_6_(i)) on our lipid bilayer systems to quantify the packing order given by 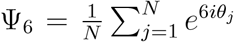, i denotes the central lipid for which the order parameter is calculated and j denotes the 6 lipids in the first solvation shell. θ_*j*_ is the angle between the central lipid i and the jth lipid with respect to a fixed global reference axis. As seen in Fig. 7(a-b), Ψ_6_ ranges from 0 (Blue) to 1 (Red) indicating degree of hexatic symmetry amongst neighbours. The parameter Ψ_6_(i) reveals the ordered-ness of lipid *i* ‘s local environment based on the neighbourhood packing symmetry. However, the overlay of Ψ_6_(i) on displacements shows poor agreement Fig. 7 (a-b) at both 288 K and 308 K.

**FIG. 7.**
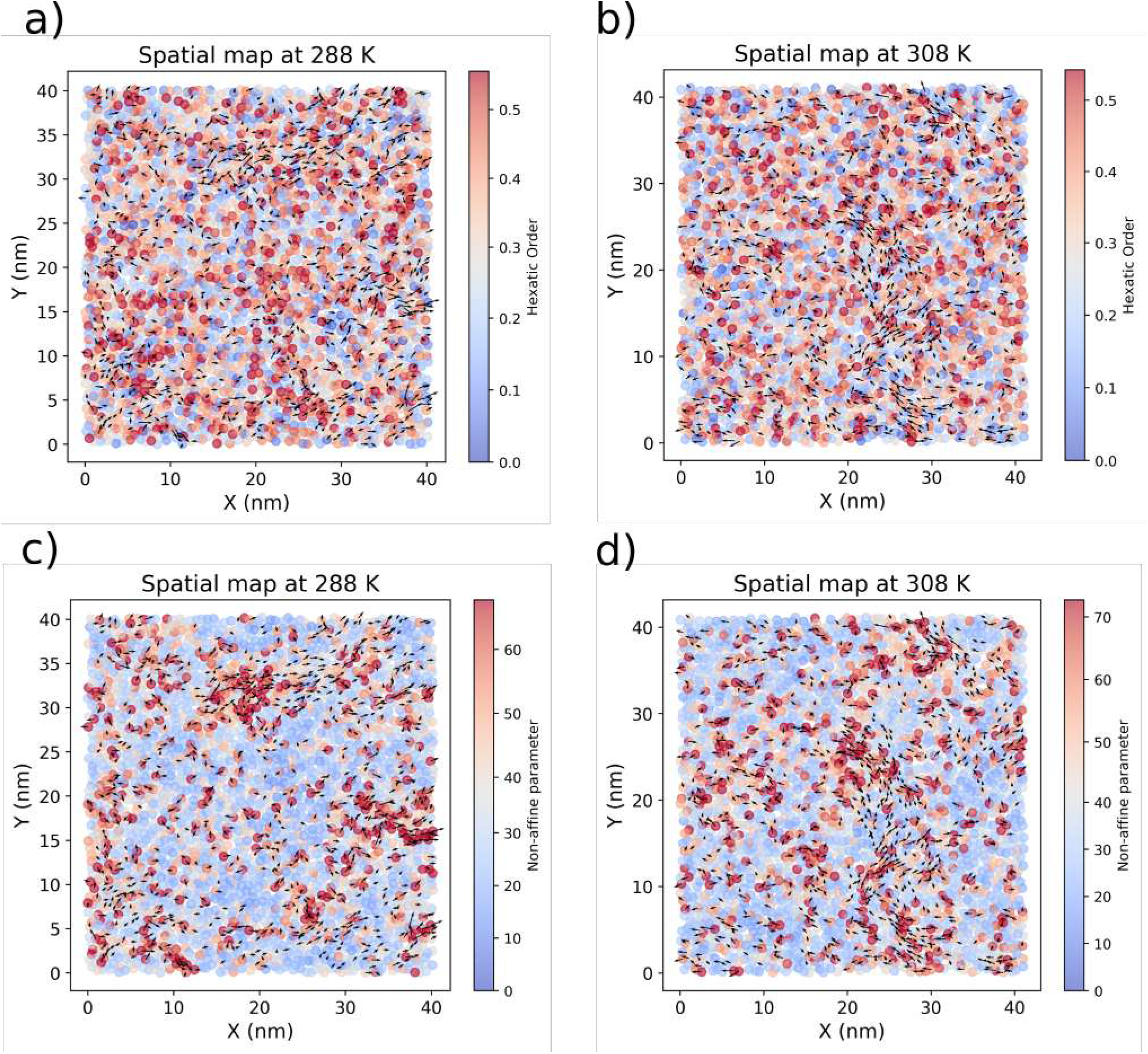
(a-b) Hexatic order parameter Ψ_6_(i) at 288 K and 308 K for each lipid *i*. Value ranges from 0 (blue) to 1 (red) (c-d) Non-affine parameter (*χ*^2^) for 288 K and 308 K. Colour coded from Blue to Red for the values

The average Ψ_6_ value is 0.3, which indicates local disorder, despite the dense packing. This parameter misrepresents the clustering behaviour due to deviations from the six-fold symmetry. To find a better characterization, we borrow a concept from the Physics of amorphous material and employ the idea of non-affine displacement (NAD) here that was formulated by Falk and Langer in 1998^34^ to identify viscoplastic deformation in non-crystalline (amorphous) material. Since dynamic heterogeneity are features of both amorphous glasses^53,54^ and lipid membranes^55–57^, we anticipated NAD to faithfully capture the spatio-temporal heterogeneities in lipid membranes. For a detailed discussion of the methodology as applied to lipid systems, we refer to a recent book chapter that we wrote in Methods in Enzymology^33^.

Mathematically we denoted NAD as χ^2^ that measure the residual non-affine part of an arbitrary deformation and is given as:

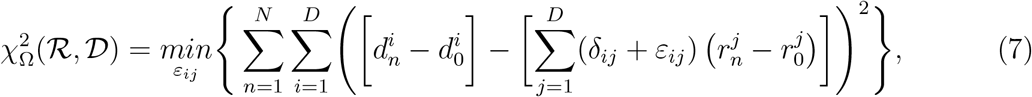

with 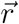 and 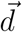 representing the coordinates of specific nodes in the reference (ℛ) and the deformed (𝒟) space respectively. i and j indicate their spatial indices in dimension D. For lipid system, we work in the 2D space. The index n runs through the N neighbouring nodes around the reference node n = 0 within the neighbourhood defined as Ω. δ_*ij*_ is the Kronecker delta function and ε_*ij*_ denotes the local strain associated with the maximum possible affine part of the arbitrary deformation, thus minimizes 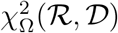, and is calculated as

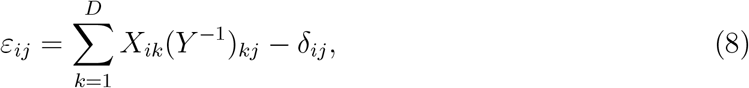

where

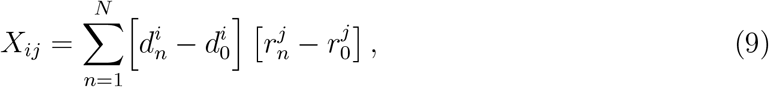

and

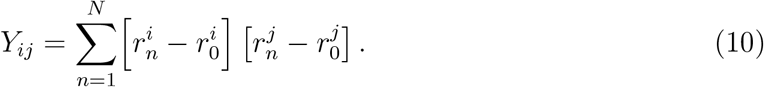

In Eq. 7, the first term in square brackets denotes the relative displacements of nodes (lipids) around the reference node (lipid) n = 0 within a neighbourhood Ω in the deformed domain. The second term in square brackets denotes the relative displacements that would have resulted if this neighbourhood was deforming under a uniform strain field ε. The quantity 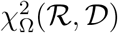 represents the mean-squared difference between these two displacements and encodes the overall change in the local topology. Consequently, for an affine deformation, 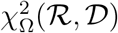 is exactly zero (see ref.^28^ for examples). A non-zero value of 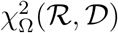 indicates the residual non-affine content of the deformation and a larger value implies a larger deviation from an affine deformation and consequently, a larger non-affine content.

With heterogeneity in the system, the lipid motions do not preserve the lattice (hexatic) structure leading to non-zero values of χ^2^. The greater the local heterogeneity around a given lipid, the greater the value of χ^2^. This makes χ^2^ an ideal quantity for characterising the time dependent local in-homogeneity present in dynamic lipid systems. Since the scores are relative, they are colour-coded here using a Yellow-Brown gradient capped at maximum of χ^2^. When overlaid (Fig.7(c-d), the spatial clustering of χ^2^ values agree quite well with the particle mobilities (Δr) close to physiological temperatures at both t_*α*_ and t_*β*_. Furthermore, the heat map also captures emergence of soft continuous channels at different timescales and temperatures accurately (see Appendix I).

Previous studies by Sodt et al. demonstrated the utility of Hidden Markov Models (HMMs) in predicting L_*o*_-L_*d*_ state of lipids using their local composition^38^. Inspired by their approach, we use χ^2^ as input parameter which is also a neighbourhood topology-based unique parameter, to identify lipids’ preference to persist as soft channels. The input parameter consists of χ^2^ (t_*β*_) scores normalised across the trajectory as emission signals. We utilise Gaussian HMM from HMMLearn python library to model the dynamic transition between two hidden states namely, soft membrane channels (relatively disordered) and strongly immobilised regions (relatively ordered). The model is trained using the Baum-Welch algorithm to determine the parameters and the most likely hidden state sequence is determined using Viterbi algorithm. The result is a transition probability matrix which indicates the lipids’ propensity to stay or alternate between the ‘ordered’ and ‘disordered’ states. We observe that the lipids increasingly prefer to exist in ordered states at 288 K (p=0.723) as opposed to higher temperatures (p=0.311 at 308 K) during short timescale fluctuations. These mixing probabilities hold true even in the diffusive regime, which is consistent with previous findings reported by Sastry and co-workers^58^.

## V. CONCLUSION

In this work, we characterised the spatial and dynamic heterogeneity of lateral membrane organisation in the liquid ordered phase. The mean squared displacement exhibits a characteristically extended sub-diffusive regime. The displacement profiles however point to inhomogeneous dynamic clustering of lipids, which is captured faithfully by the NAD (χ^2^) order parameter. Though subtle, we observe signatures of dynamic heterogeneity through overlap function and dynamic susceptibility and also extract the timescales associated with the spatial segregation. The extensive deviation from exponential decay in Q(t) also points to the existence of multiple time-scales within the system. The particle distribution evolution through van-Hove function confirms the presence of short multiple spatially correlated segments of lipids in displacement profile. Interestingly, the structural relaxation dynamics is Arrhenius-like indicating that the L_*o*_ phase has the properties of a strong glass former. This is analogous to previous findings on soft colloidal properties^59^ of a crowded cytoplasm in living cells^60–62^, further highlighting the importance of strong glass-forming features in biology. However unlike previous studies, the glassy behaviour in membranes appears to be intrinsically driven by thermal fluctuations, requiring no external energy inputs^2,63^. This implies that membranes naturally resist drastic temperature-dependent structural re-organisation, while still supporting surface level molecular diffusion dynamics.

## Supporting information

Gif movie files of lipid dynamics at different temperatures

## APPENDIX I: ROBUSTNESS OF THE NON-AFFINE DISPLACEMENT PARAMETER

In the previous sections, we analysed the heterogeneity via a variety of markers, along with making visual observations. However, till date we have not been able to connect such markers to the diffusion of lipids in a robust manner. In this section, we consider a theoretical model which explains a possible mechanism for non-Gaussian diffusion to occur. To describe anomalous diffusion (Brownian but non-Gaussian), Chechkin and co-workers^20^ introduced a minimal model where the particle position 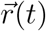 follows a set of stochastic equations of motion given by:

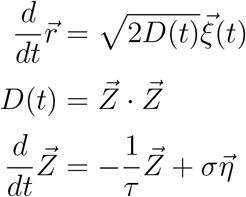

Considering the diffusion coefficient D(t) as random in time, they express D(t) in terms of the square of the Ornstein-Uhlenbeck process 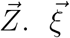 and 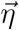 are normally distributed white Gaussian noise. The analogy in these set of equations is that 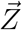 drives the diffusion of the particle, by determining the local time-dependent diffusion constant. 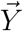on the other hand also follows a relaxation time type equation, with a random noise source which can sometimes give it large values. This set of stochastic equations can be solved to show that 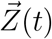 follows a normal Gaussian distribution, and as a result 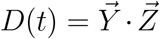 is a χ^2^ distribution with n degrees of freedom. However, notice that χ^2^ is the error that arises from the best fit calculation for the ϵ_*ij*_ tensor. Such an error is also a χ^2^ distribution, albeit with a different degree of freedom. This motivated us to make an identification of D(t) with the χ^2^ non affine parameter, and forms the primary basis of this analysis. By finding an analogy between the χ^2^ value and the diffusion rate of the particles, we demonstrate an unequivocal relationship between χ^2^ as a marker for lipid diffusivity. Since diffusion is proportional to standard deviation of displacement vector components σ_Δ*r*_, we evaluate it as a function of local order (χ^2^) and report in Fig. 8.

**FIG. 8.**
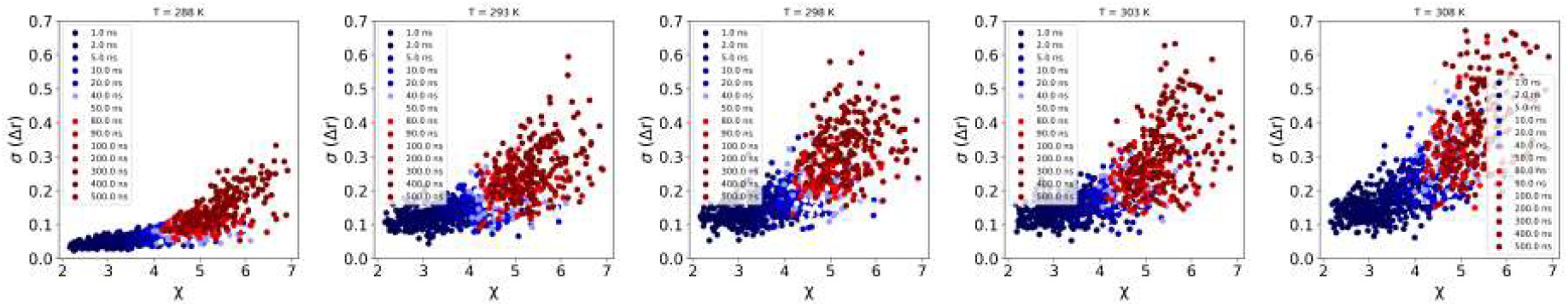
The *σ*(Δ*r*) vs *χ* plot has a growing relationship that is consistent across temperatures.

## ACKNOWLEDGEMENT

HS thanks the Ministry of Education, Government of India, for the graduate fellowship. AS acknowledges the financial support from the Indian Institute of Science-Bangalore and the high-performance computing facility “Beagle” that was set up from grants by a partnership between the Department of Biotechnology of India and the Indian Institute of Science (IISc-DBT partnership programme). AS thanks the DST for the National Supercomputing Mission grants (DST/NSM/R&D-HPC-Applications/2021/03.10, DST/NSM/R&D-HPC-Applications/Extension Grant/2023/27). FIST program sponsored by the Department of Science and Technology and UGC, Centre for Advanced Studies and Ministry of Human Resource Development, India. AS would also like to thank the Teams Science Grant from the DBT-Wellcome Trust India Alliance (Grant number: IA/TSG/21/1/600245). AS also thanks the DBT National Network Project (NNP) grant (BT/PR40323/BTIS/137/78/2023) and the Matrics grants (MTR/2023/001040) from the Science and Engineering Board (SERB), India. EL was supported by NSF award number MCB-2121854.

## AUTHOR CONTRIBUTIONS

AS conceived the idea behind the project. SSI, HS and AS designed the research. HS wrote the pre and post-processing Python tools needed for the analyses. HS performed the research, generated the data and carried out the first set of analyses. SSI, AB, EL and AS participated in the analysis of the data and proposed advanced analyses ideas. AS supervised the study. HS prepared the first draft of the paper, and all others polished it together.

## CONFLICT OF INTEREST

The authors declare no potential conflict of interest.

## DATA AVAILABILITY STATEMENT

Our codes, models, and curated datasets are publicly available at the Figshare repository https://figshare.com/s/3adebead8c5785323449.

## Notes

https://github.com/codesrivastavalab/glassyDynamicsLipids

https://figshare.com/s/3adebead8c5785323449

### Competing Interest Statement

The authors have declared no competing interest.

