## Supplementary figures and images for "Signatures of glassy dynamics in highly ordered lipid bilayers with emergence of soft dynamic channels"

### T288-Relaxation.gif

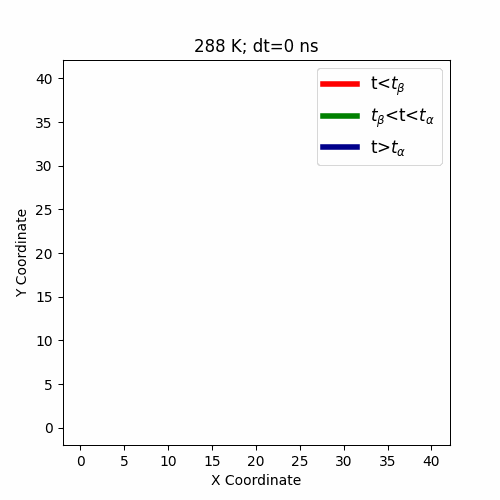

### T293-Relaxation.gif

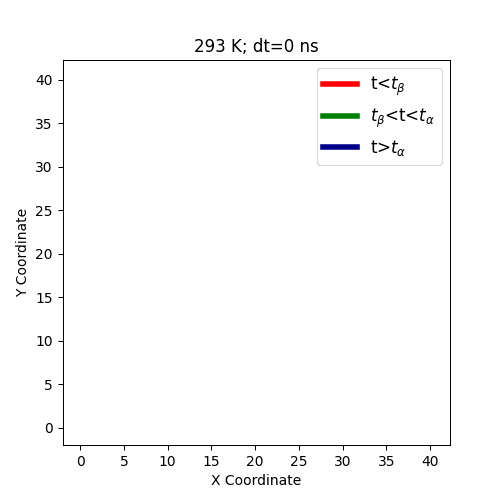

### T298-Relaxation.gif

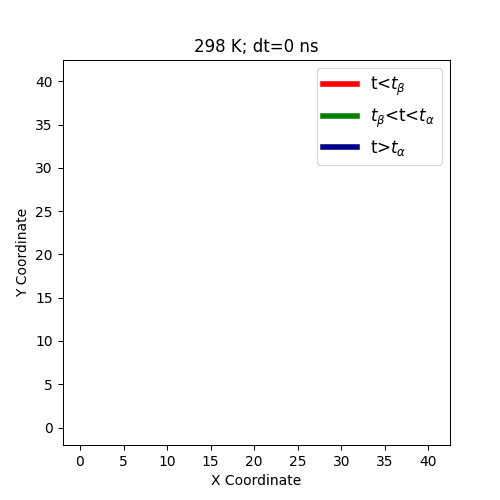

### T303-Relaxation.gif

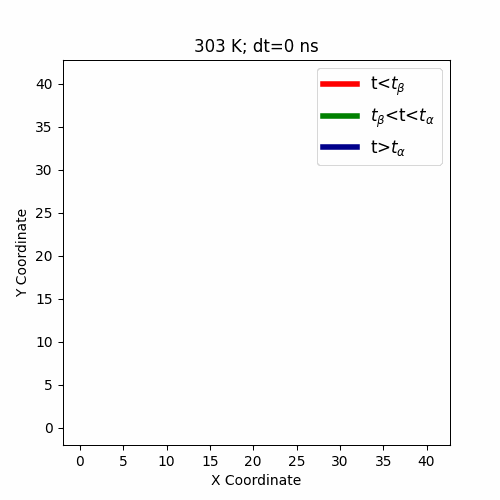

### T308-Relaxation.gif

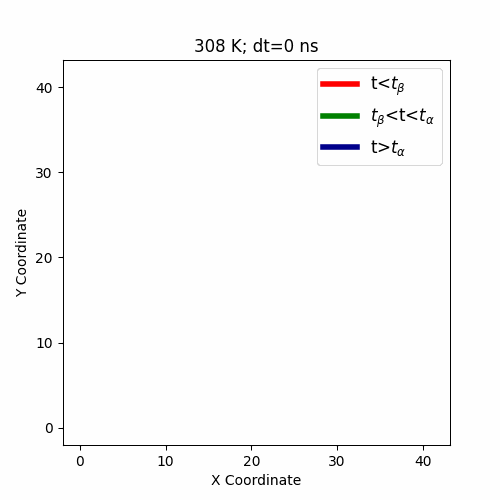
